# Sourdoughs fermented by autochthonous *Lactobacillus* strains improve the quality of gluten-free bread

**DOI:** 10.1101/2020.08.21.260653

**Authors:** Yousef Nami, Mehdi Gharekhani, Mehran Aalami, Mohammad Amin Hejazi

**Author notes:** **Corresponding author:** Mohammad Amin Hejazi; Address: Agricultural Biotechnology Research Institute of Iran, Daneshghah Street, Tabriz, Iran. P. O. Box: 5156915598.

## Abstract

Sourdoughs based on fermentation by lactobacilli have the potential to produce gluten-free maize-based bread with acceptable technological and rheological characteristics, nutritional quality and a more prolonged shelf life. Of the 17 treatments compared (with or without sourdough, and involving single and multiple LAB species), treatments 12C (*Lactobacillus brevis, L. sanfranciscensis* + *L. plantarum*) and 8C (*L. brevis* + *L. paralimentarius*) showed the lowest rate of complex modulus, while treatments 11C (*L. sanfranciscensis* + *L. brevis* + *L. paralimentarius*) and 2C (*L. brevis*) led to the greatest reduction in baking loss. The crumb moisture content of all of the formulations decreased with storage. Breads produced with treatment 2C (*L. brevis*) had the highest crumb moisture content when freshly baked; while loaves produced with treatment 3C (*L. paralimentarius*) had the highest crumb moisture content after four days of storage. A sensory evaluation indicated that sourdough-based maize breads were superior to both control and chemically acidified breads. The optimal treatments were to use sourdough seeded with treatment 2C (*L. brevis*), with treatment 4C (*L. plantarum*), with treatment 8C (*L. brevis* + *L. paralimentarius*) or with treatment 11C (*L. sanfranciscensis* + *L. brevis* + *L. paralimentarius*).

**Significance Statement:** Of the 17 treatments compared, treatments 12C and 8C showed the lowest rate of complex modulus, while treatments 11C and 2C led to the greatest reduction in baking loss. The optimal treatments were to use sourdough seeded with treatment 2C, with treatment 4C, with treatment 8C or with treatment 11C.

## 1. Introduction

Bread is the most important food for most people, especially in developing countries. Bread is usually made from wheat flour, but bread from rye, barley, and millet is also common. On average, about 60-65% of calories and protein and 2-3 gr of mineral salts are provided daily by eating bread. Bread has always been one of the cheapest sources of energy and protein for human. Bread has a prominent role in providing dietary fiber, certain minerals such as calcium, iron, group B vitamins as well as vitamin E. Since the products of the fortune industry have a special place in the food basket of the community, even celiac patients, the use of gluten-free flours such as rice, corn, sorghum, cassava, amaranth, and quinoa is inevitable (Kumar et al., 2019;Skodje et al., 2019).

Gluten-free (GF) cereal-based foods are required for patients suffering from coeliac disease (Matos and Rosell, 2015). Producing GF bread acceptable to the consumer is difficult, largely because gluten is the basis of the viscoelastic network required to create bread’s characteristic spongy texture. In addition, GF bread tends to suffer from poor color, a short shelf-life and a generally unsatisfactory organoleptic score (Rinaldi et al., 2017). Although the formulation of the dough used to produce GF bread is critical, the fermentation and baking conditions also influence the quality of the product.

Lactic acid bacteria (LAB) can be used to acidify doughs (De Vuyst and Neysens, 2005), a process which improves the leavening process and has a positive effect on the quality and shelf-life of wheat bread (Scarnato et al., 2017). Sourdoughs based on a combination of LAB and yeast not only extend the shelf-life of the bread but also improve its nutritional value, flavor and aroma (Poutanen et al., 2009). The use of LAB does, however, risk-averse effects on dough rheology, since some strains exhibit proteolytic activity. Because of the sensitivity of dough rheology to any ingredients and its related consequence on the GF bread quality, screening and introduction of new strains of LAB for application in GF bread production is of importance.

One of the most commonly used LAB strains in the production of GF bread is *Lactobacillus sanfranciscensis* (Gänzle et al., 2008). Both (Scarnato et al., 2017) and (Vernocchi et al., 2008) have used this LAB, in conjunction with *Candida milleri,* to improve the aroma and lengthen the shelf-life of a range of GF products.

The present study aimed to improve the quality of maize-based bread through the use of different native LAB isolates belonging to species, namely *L. sanfranciscensis, L. plantarum, L. brevis,* and *L. paralimentarius*.

## 2. Materials and methods

### 2.1. Bacteria isolation and identification

Five lactic acid bacteria (LAB), which were isolated from traditional sourdough of East Azerbaijan, were obtained from the bacterial collection of Agricultural Biotechnology Research Institute of Iran (ABRII). These bacteria were inoculated in De Man Rogosa Sharpe medium (MRS) under sterile conditions and incubated at 37 °C for 24 h. For molecular identification of bacteria, 16s-rRNA fragments were amplified according to a method of (Haghshenas et al., 2017). The samples were identified and characterized at species levels using BLAST software (http://blast.ncbi.nlm.nih.gov/Blast.cgi) and by comparing them with the deposited sequences in NCBI and GenBank.

### 2.2. Preparation and characterization of sourdoughs

Sourdoughs were fermented using one, or a combination of the four LAB species *L. sanfranciscensis, L. brevis, L. paralimentarius,* and *L. plantarum.* In addition, chemically acidified (CA) and control sample (without starter) were used for fermentation of sourdoughs. In total, 17 treatments were compared, as stated in table 1. The doughs were made by mixing maize flour and water to obtain a dough yield (DY) of 200. The various sourdoughs were prepared by seeding maize dough with selected starter cultures. Fermentation was carried out at 30°C for 24 h (Nami et al., 2019), after which the following parameters were measured, following the methods given by Paramithiotis et al. (2010): pH, total titratable acidity, hydrogen peroxide content, and diacetyl value. A count of the bacteria was also performed.

**Table 1:** Signs and abbreviations used instead of treatments titles in sourdough, dough and bread and formulation of GF bread dough preparation

| Signs and abbreviations of used strains in treatments |  | formulation of GF bread dough preparation |  |  |  |
| --- | --- | --- | --- | --- | --- |
| Treatment s | LAB Strains | Raw materials | Control Dough | Sourdough Dough | Acid-Chemical Dough |
| 1C | *L. sanfranciscensis* | Flour | 100 | 92.5 | 100 |
| 2C | *L. brevis* | Salt | 2 | 2 | 2 |
| 3C | *L. paralimentarius* | Sugar | 5 | 5 | 5 |
| 4C | *L. plantarum* | Yeast | 3 | 3 | 3 |
| 5C | *L. sanfranciscensis* + *L. brevis* | Egg | 12 | 12 | 12 |
| 6C | *L. sanfranciscensis*+ *L. paralimentarius* | Sodium Caseinate | 1.5 | 1.5 | 1.5 |
| 7C | *L. sanfranciscensis*+ *L. plantarum* | Skim milk | 5 | 5 | 5 |
| 8C | *L. brevis*+ *L. paralimentarius* | Guar Gum | 3 | 3 | 3 |
| 9C | *L. brevis*+ *L. plantarum* | Edible Oil | 5 | 5 | 5 |
| 10C | *L. paralimentarius*+ *L. plantarum* | Sourdough | - | 15 | - |
| 11C | *L. sanfranciscensis*+ *L. brevis*+ *L. paralimentarius* | Acid* | - | - | 0.09 |
| 12C | *L. sanfranciscensis*+ *L. brevis*+ *L. plantarum* | Water | 110 | 102.5 | 110 |
| 13C | *L. brevis*+ *L. paralimentarius*+ *L. plantarum* |  |  |  |  |
| 14C | *L. sanfranciscensis*+ *L. paralimentarius*+ *L. plantarum* |  |  |  |  |
| 15C | *L. sanfranciscensis*+ *L. brevis*+ *L. paralimentarius*+ *L. plantarum* |  |  |  |  |
| 16C | Chemically acidified (CA) sample |  |  |  |  |
| 17C | Control (CO) sample (without starter) |  |  |  |  |

\* Mixing lactic acid - acetic acid in a ratio of 4: 1 (V.V)

### 2.3. Dough rheology

Standard dough rheology tests were conducted, following (Moroni et al., 2011). The doughs were held at a constant temperature of 30°C, using a Peltier Plate System attached to a water circulation unit. Briefly, controlled stress and strain rheometer (Antoon Paar MCR 301, Ostfildern, Germany) was used to measure rheological parameters. Parallel plate geometry was used for measuring dough samples. Dough samples were allowed to rest for 10 min prior to evaluation and doughs were incubated for one hour at fermentation conditions (30 °C and 75% humidity) before analysis.

### 2.4. GF bread preparation

The GF bread recipe comprised maize flour, water, sugar, salt, egg, baker’s yeast, skimmed milk, sodium caseinate, guar gum, edible oil and sourdough (the amount of each one is stated in table 2); this formulation produced ~500 g of dough with a DY of 200. A 150 g portion of each dough was baked in a tin (15 cm × 8.5 cm × 5.7 cm) at 225°C for 30 min. The loaves were stored in polyethylene bags after cooling to room temperature (Moore et al., 2008).

**Table 2:**
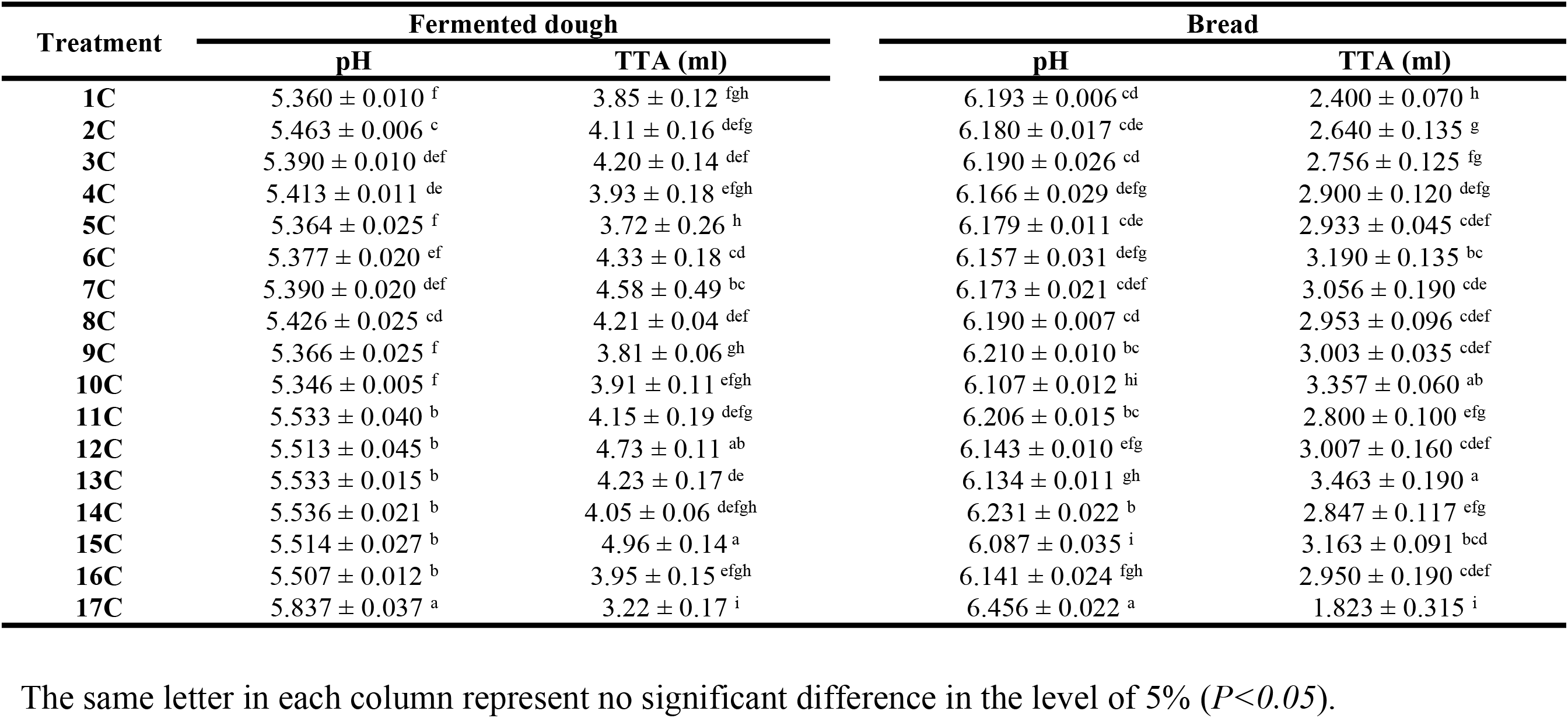
Changes in pH and TTA in the dough and gluten-free bread

### 2.5. Bread and dough Physico-chemical attributes

#### 2.4.1. pH and total titratable acidity

The pH of the dough and bread was obtained by soaking a 10 g sample in 90 mL distilled water and measuring the pH with a standard pH meter. Total titratable acidity values were obtained by recording the volume of 0.1 M NaOH needed to raise the same samples’ pH to 8.5.

#### 2.4.2. Diacetyl and hydrogen peroxide production

Diacetyl production was calculated by mixing 10 g sourdough samples in 90 mL distilled water. Afterward, 7.5 mL of hydroxylamine solution (1 M) was added to 25 ml of the homogenized mixture and samples were titrated by 0.1 N HCl to final pH 3.4. The equivalence factor of HCl to diacetyl is 21.52 mg. The concentration of produced diacetyl was measured according to a method of (Edema and Sanni, 2008).

Hydrogen peroxide production was evaluated by adding 25 mL of 10% H_2_SO_4_ to 25 mL of homogenized mixture (from the same batch used for diacetyl). It was then titrated with 0.1 N potassium permanganate (KMnO4) so that the pale pink color persisted for 15 s before de-colorization. Each mL of 0.1 N KmnO_4_ is equivalent to 1.701 mg of H_2_O_2_. The concentration of produced H_2_O_2_ was calculated as follows (Edema and Sanni, 2008):

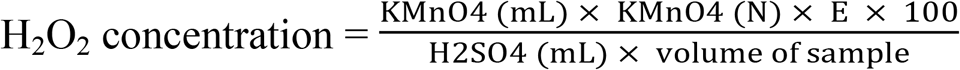

#### 2.4.3. LAB cell counts

A 10 g sample of sourdough was homogenized in 90 mL 0.15 M NaCl and serial dilutions were prepared in phosphate buffer saline (PBS). The dilutions were plated in triplicate on MRS agar and incubated for 48 h at 30°C.

#### 2.4.4. Crumb and crust color

Bread crumb and crust color evaluations were performed following (Marti et al., 2017). Values for L*, a*, and b* (as measures of lightness, redness-greenness, and yellowness–blueness, respectively) were measured for each sample. Each measurement was replicated three times.

#### 2.4.5. Specific bulk volume and height

The specific volume of three replicate loaves per formulation was measured using the rapeseed displacement method (AACC10-05), performed one hour after baking. The loaves were weighed and their specific volume was determined from the volume/mass ratio (mL g^-1^). A pair of digital calipers was used to estimate loaf height.

#### 2.4.6. Moisture content

The moisture content of the breads was measured following AACC standard method 44-16 (AACC, 2000). The moisture content of the crust and central crumb of fresh loaves was recorded, and similar measurements were taken after storage of the loaves for two and four days.

#### 2.4.7. Baking loss

Baking loss was obtained from weight measurements taken before and after baking, according to the following formula:

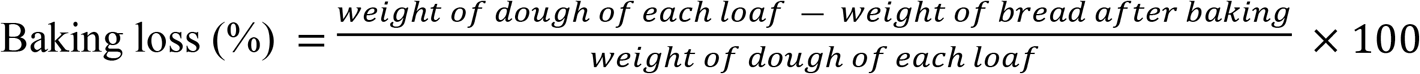

#### 2.4.8. Bread porosity

The (Wolter et al., 2014) method was used to obtain estimates of the percentage crumb porosity.

#### 2.4.9. Crumb and crust hardness

A textural profile analysis was performed on the crumb and a penetration test used quantifies crust hardness. Both tests utilized methods described by (Crowley et al., 2002).

### 2.5. Organoleptic attributes of bread

The organoleptic attributes of the breads were determined following a standard protocol (ISO 8587, 1988) which employed a panel of twelve trained judges. An overall organoleptic score was based on the individual assessments of crust and crumb color, porosity, elasticity, acidic smell, texture softness, chewiness, and taste. The texture characteristics, chewiness, and taste were first evaluated 2 h after baking, then again after two and four days of storage.

### 2.6. Shelf-life evaluation

The breads were enclosed in polyethylene bags after cooling and cutting with a sterile knife. The number of days of storage at room temperature required for the appearance of mold was considered as the bread’s shelf life (Moore et al., 2008).

### 2.7. Statistical analyses

The data were statistically analyzed using routines implemented in SAS v9.0 software (SAS Institute, Cary, NC, USA). The data are presented in the form mean ± standard error (*n=*3). The significance threshold adopted for the analyses of variance was 0.05.

## 3. Results and Discussion

### 3.1. Bacteria isolation and identification

The fragments (1500 bp) of the 16S-rRNA gene were sequenced for molecular identification of the isolates. Based on sequencing results, five strains belonged to four species, namely *Lactobacillus brevis, Lactobacillus sanfranciscensis, Lactobacillus plantarum and Lactobacillus paralimentarius* (Table 1).

### 3.2. Sourdough preparation and characterization

Both the single LAB strain and combined strain starter cultures significantly increased the acidity (decreased the pH) and total titratable acidity of the sourdoughs. The highest total titratable acidity (9.05 ± 0.095 mL) was associated with the treatment 9C (*L. brevis* + *L. plantarum*) and the lowest (6.24 ± 0.360 mL) with the control doughs to which no starter had been added. The sourdough which accumulated the most diacetyl (34.45 ± 1.510 mg mL^-1^) was produced using treatment 6C (*L. paralimentarius* + *L. sanfranciscensis*), while the one accumulating the most hydrogen peroxide (3.90 ± 0.126 mmol L^-1^) was produced using treatment 1C (*L. sanfranciscensis*). The variation in the size of the LAB populations at the end of the fermentation period is illustrated in Fig. 1. The largest population developed from treatment 5C (*L. sanfranciscensis* + *L. brevis*). An analysis of variance confirmed that the nature of the starter culture had a significant effect on the growth of the LAB in the sourdough.

**Figure 1:**
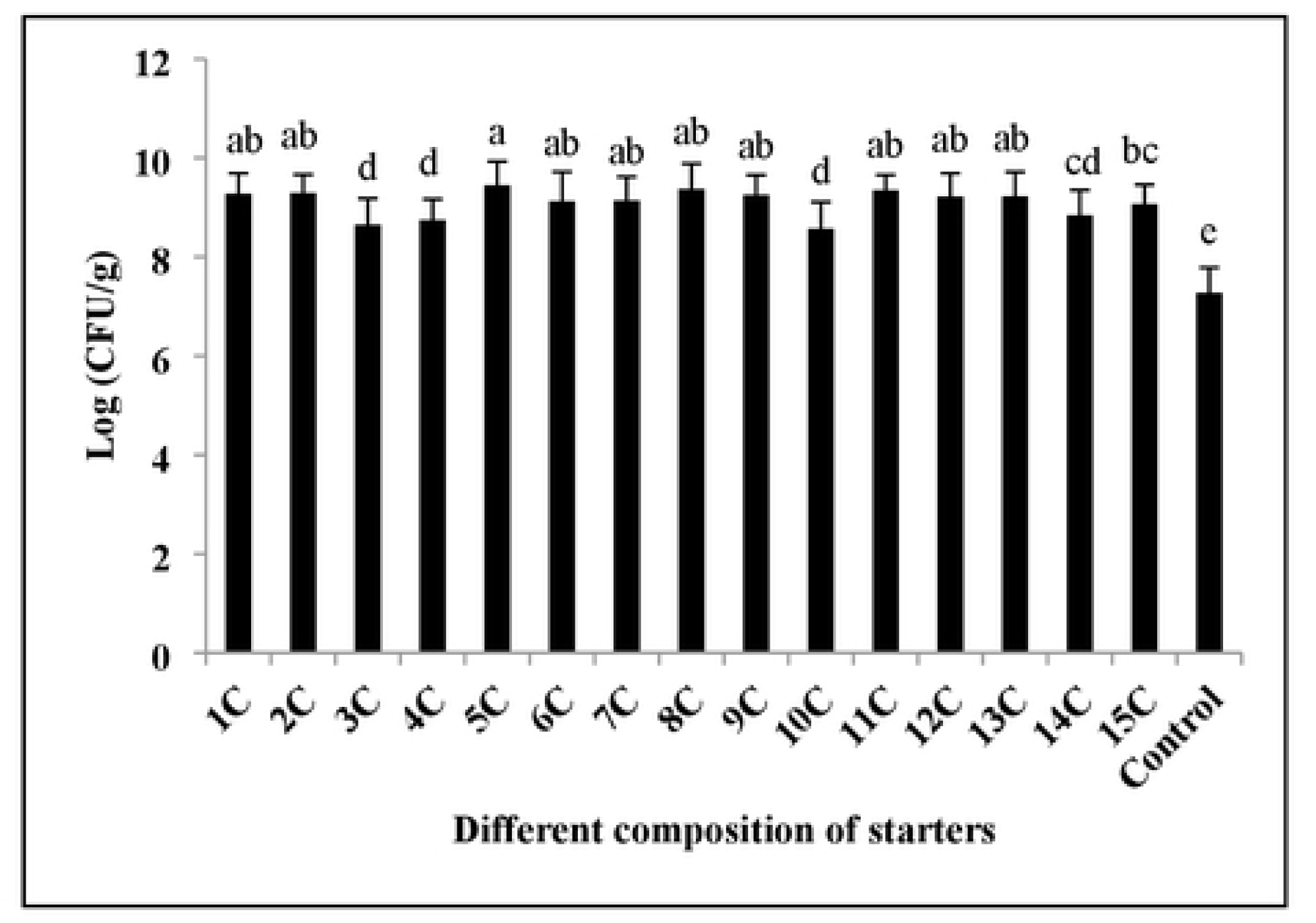
The impact of different starters on the LAB population of maize sourdough.

The effect of the various starter cultures on dough and bread pH and total titratable acidity (Table 3) implied that the incorporation of sourdough significantly reduced pH and increased acidity in both bread and sourdough. Treatment 15C (sourdough seeded with *L. plantarum* + *L. brevis* + *L. paralimentarius* + *L. sanfranciscensis*) was associated with the lowest bread pH, while treatment 13C produced the highest total titratable acidity.

**Table 3:**
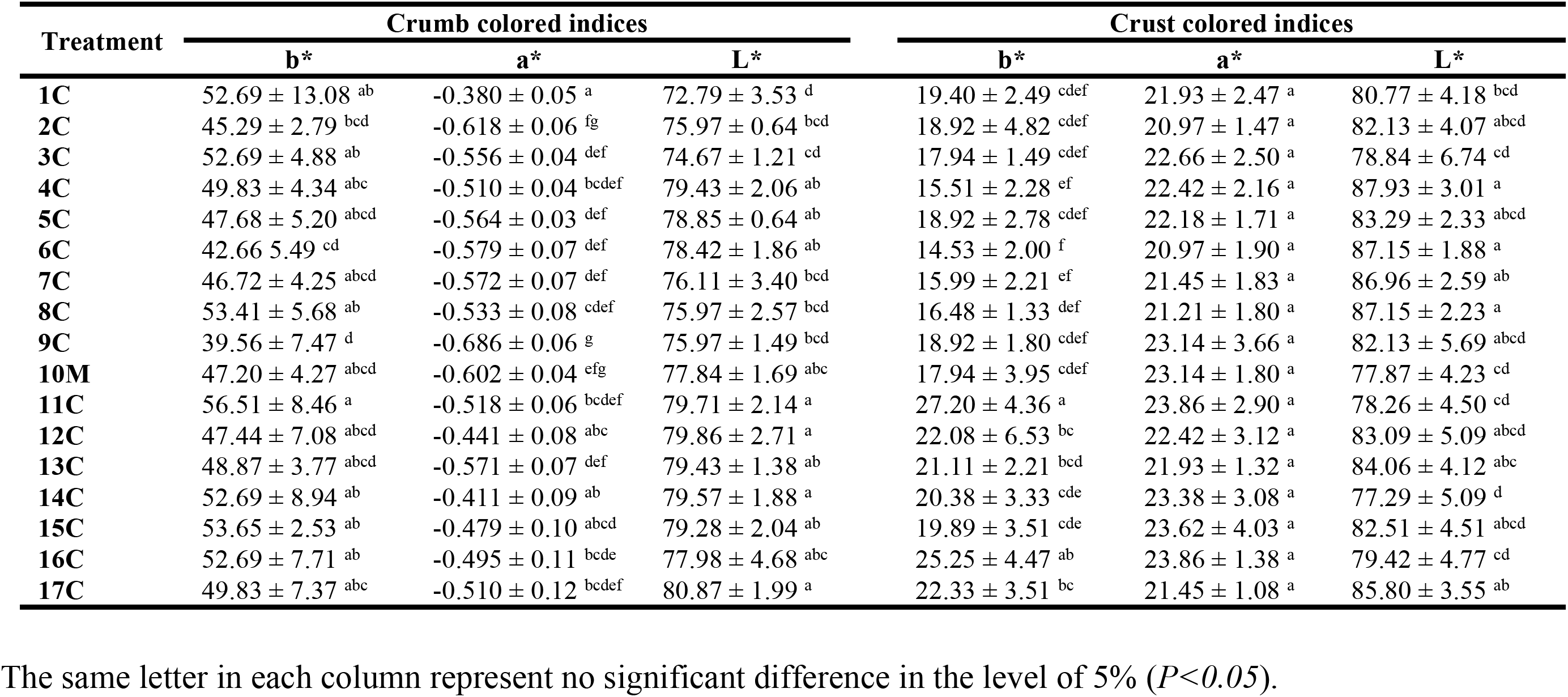
The effect of different starters on colored indices of maize bread crumb and crust

Effects of starters on the amount of diacetyl and hydrogen peroxide production was statistically significant (P<0.05) in sourdoughs (not shown). Starter-containing sourdoughs showed a higher amount of diacetyl and hydrogen peroxide than starter-free sourdoughs. Among the sourdoughs, starter 6C showed a significant increase in the amount of diacetyl comparing to other starters, while starter 1C showed the highest amount of hydrogen peroxide.

### 3.2. Dough rheology

A frequency sweep test was performed to evaluate the effect of the nature of the starter culture on the rheological properties of control (no additives, CO) and chemically acidified (CA) breads. As shown in Fig. 2, the G* parameter increased with ω throughout and was highest in CO dough. Both the addition of acid to the dough and the inclusion of sourdough reduced dough stiffness. The comparison between CA dough and those produced by seeding with a sourdough starter cultures revealed that the latter induced a greater fall in G*. Treatments 2C, 7C, 8C, 13C, and 15C all produced doughs with a lower G* than those formed by seeding with any of the other starter cultures. For all doughs, the δ value decreased with increasing ω. CO dough was the most elastic, followed by CA dough; doughs produced from treatments 8C and 2C were associated with the highest δ value.

**Figure 2:**
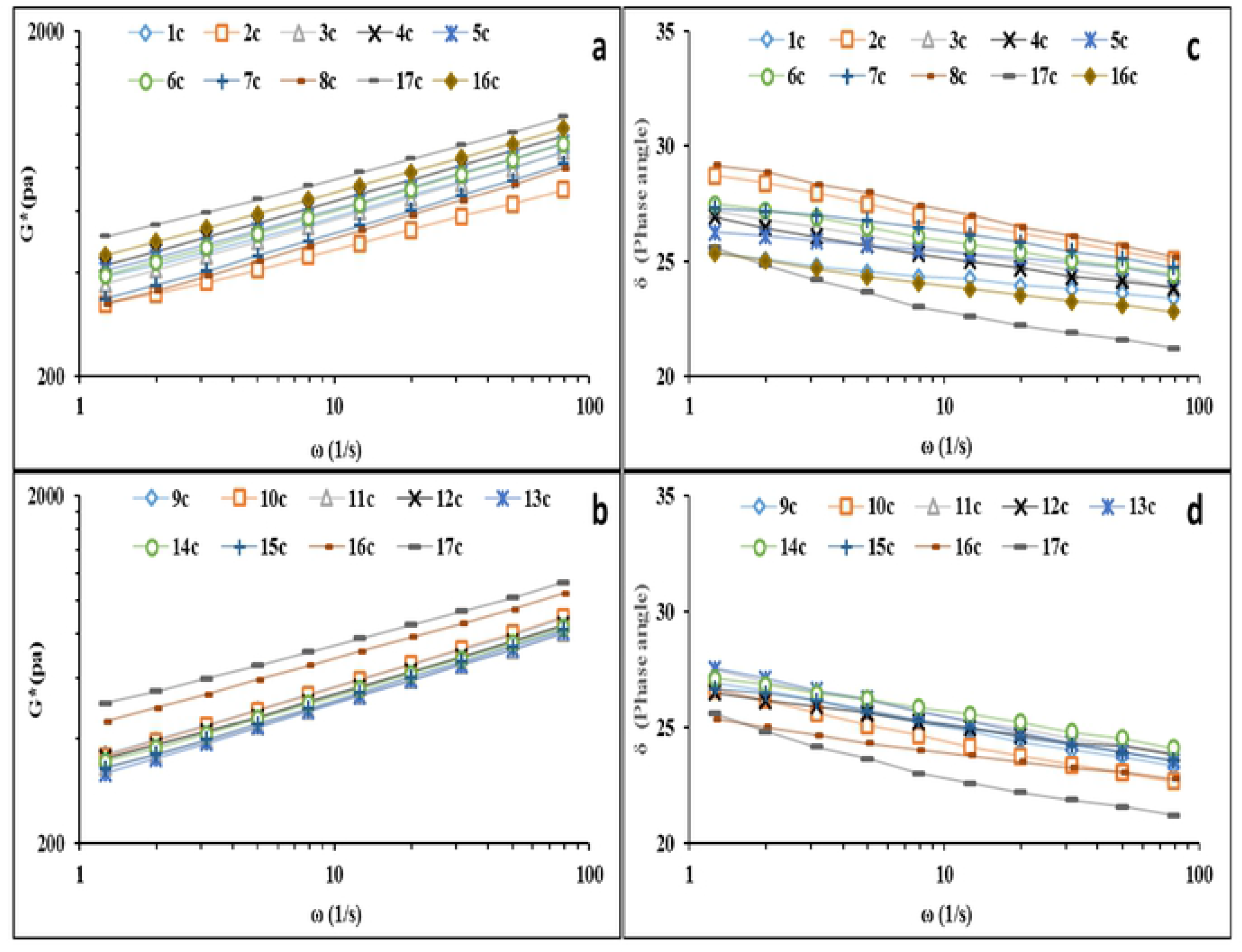
Comparison of the complex modulus (G*, Panels a, b) and phase angle (δ, Panels c, d) of maize doughs with increasing frequency (ω).

(Edema, 2011) have reported that employing *L. brevis* in a starter culture for maize sourdough was highly effective in terms of boosting its diacetyl content, while the best results with respect to hydrogen peroxide content were obtained using a combination of *L. plantarum* and *L. brevis*. These conclusions were borne out in the present experiments. According to (Clarke et al., 2002), in wheat breads, the addition of sourdough (*L. plantarum* and *L. brevis*) had the effect of increasing δ and reducing dough elasticity. In bread formulated with GF flour, the G* value is boosted by the inclusion of sourdough, thereby stiffening the dough.

### 3.3. Bread crumb and crust color

Colour, texture, and aroma are all important quality traits in bakery products (Esteller et al., 2006). Colour is usually quantified by a combination of the parameters L*, a* and b*. The effect of the various sourdoughs on bread crumb and crust color is shown in Table 4. The addition of sourdough decreased light crumb compared to that present in CO bread. Treatment 1C (*L. sanfranciscensis*) induced lower light and the most intense crumb redness. There was no significant difference between sourdough starter cultures and CO with respect to either L* or a*, but the use of most of the various sourdoughs did decrease b* (the yellowness crust value) - the exceptions were treatments 11C, 12C (*L. sanfranciscensis+ L. brevis+ L. plantarum*) and 13C.

(Aplevicz et al., 2014) showed that the crust color of sourdough-based breads (*L. plantarum*) was lighter than starter-based breads, but was less bright than that of CO bread.

**Table 4:**
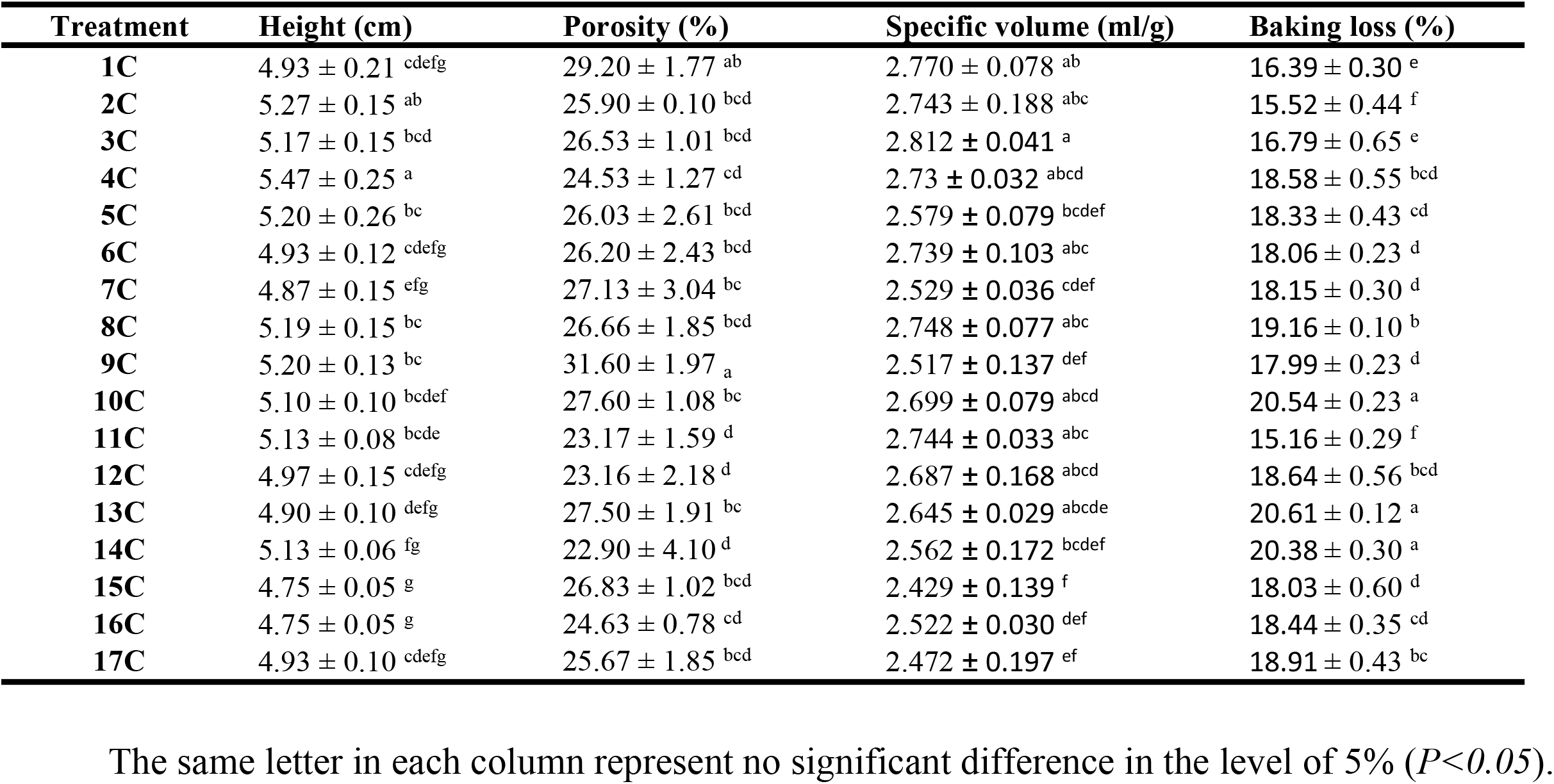
The effect of different starters on traits of gluten-free maize bread

### 3.4. Crumb and crust hardness

Crust and crumb hardness was significantly affected by both the identity of the starter culture type and the storage time. Breads produced from doughs subjected to treatment 2C produced the lowest crumb hardness (800.05 g) measured on fresh loaves, but in general, the sourdough-based breads produced a harder crumb structure than did either the CO or the CA breads. The highest crumb hardness values for fresh loaves were associated with breads prepared from doughs subjected to treatment 13C. After storage for two days, breads produced from doughs subjected to treatments 2C, 8C and 7C had a softer crumb than the others, while after four days of storage, breads produced from doughs subjected to treatments 8C (1273.71 g) and 7C (1227.44 g) had a softer crumb than those prepared from sourdoughs seeded with a single LAB starter culture and CO bread. Crust hardness declined during storage (data not shown). Breads produced from doughs subjected to treatment 8C had a hardness value of 474.46 g when fresh and 401.44 g after two days: this treatment produced the softest crust. The hardest crusts were associated with CA and CO breads. Treatment 15C produced loaves with the softest crust at the end of the storage period. The hardest crusts were associated with breads produced from doughs subjected to treatments 11C, 5C, 9C, 14C, along with CO and CA breads.

With respect to the breakdown of the texture of bread during storage, the reductions observed in the maize-based breads are consistent with the literature (Clarke et al., 2002). According to Moore et al. (2008), the texture of GF breads is influenced by storage time and its interaction with the formulation of dough: crumb hardness increases over time, but more so in CA than in either sourdough-based or CO bread. The inclusion of sourdough has been documented to delay the staling of GF breads (Corsetti et al., 2000) have suggested that LAB-mediated acidification encourages starch hydrolysis, proteolysis, and various other physicochemical changes during the course of storage. When (Moroni et al., 2011) evaluated buckwheat-based sourdoughs seeded with different starter cultures for the production of wheat breads; it was found that their inclusion had a marked effect on dough rheology: it reinforced the action of the gluten network, and so reduced dough elasticity. This produced both an increase in loaf specific volume and a softening of the crumb structure. The rate at which the crumb hardens is influenced both by the starter culture type and the proportion of the dough made up by sourdough (Novotni et al., 2013): at low proportions of the latter, there was no effect of adding the sourdough, but higher proportions (22.5% and 30%) were effective in increasing crumb firmness.

### 3.5. Specific volume and height of breads

The starter cultures had a significant effect on both the height and specific volume of the bread. The highest specific volume was produced by breads subjected to treatment 3C (sourdough seeded with *L. paralimentarius*); the lowest specific volume was associated with CA bread. According to (Mert et al., 2014), GF doughs can be softened by the addition of sourdough, as this encourages the expansion of gas bubbles during fermentation; it also raises the specific volume of the loaf since it improves the dough’s capacity to retain carbon dioxide.

### 3.6. Moisture content of breads

Analyses of variance (not shown) suggested that the nature of the starter culture, the post-baking storage time and their interaction all had a significant effect on the moisture content of the crust and crumb. The moisture content of the crumb was consistently reduced as the storage time was extended, while that of the crust increased. Immediately after baking, bread made from dough subjected to either of the treatments 12C and 11C retained the most moisture; after four days of storage, breads made from dough subjected to either of the treatments 3C, 7C, 9C and 10C (*L. paralimentarius+ L. plantarum*) retained the most moisture. The lowest moisture content after storage was recorded by CA loaves. In conjunction with the maize crust moisture, bread made from dough subjected to either of the treatments 7C and 8C retained the least moisture immediately both after baking and after four days of storage.

The moisture content of sourdough-based wheat breads falls during storage (Aplevicz et al. 2014), although breads based on sourdoughs formulated with *L. plantarum* are able to retain higher moisture content and thus are more palatable than CO breads. As a result, our findings are consistent with these researchers. However, (Ryan et al., 2011) were not able to find any difference in the crumb moisture content of CO, CA and sourdough-based bread fermented with *L. amylovorus.* There was, similarly, in the materials investigated by (Barber et al., 1992), no LAB-dependent impact on moisture content following either baking or storage time. The high moisture content of GF breads reflects the need to include much more water in the dough than is necessary in wheat bread formulations. For example, the oat-based breads described by (Hüttner et al., 2010) had a moisture content of 58-61%. According to Tamani et al. (2013), a benefit of including sourdough based on *L. delbrueckii* and *L. helveticus* is that the crumb moisture content falls less rapidly over time. When bread’s moisture content is reduced, cross-linking between starch and protein accelerates in intensity, leading to increasing stiffness (Symons and Brennan, 2004). A high content of exopolysaccharides in sourdough may be responsible for the improvement in water retention and hence a softer crumb structure. However, it may that the qualitative nature of the exopolysaccharides present is as important as their quantity. According to (Wolter et al., 2014), the addition of sourdough containing *L. plantarum* decreased baking loss in GF breads, which was also the case in the present study.

### 3.7. Baking loss

The choice of starter culture had a significant effect on baking loss (Table 4). Doughs subjected to treatment 11C showed the lowest percentage of weight loss after baking, while the highest baking losses were observed following treatments 10C, 13C, and 14C (*L. sanfranciscensis+ L. paralimentarius+ L. plantarum).*

### 3.8. Bread porosity

The nature of the sourdough significantly influenced bread porosity (Table 4), with treatments 9C and 14C showing, respectively, the highest and the lowest porosity.

The porosity of quinoa-based CO bread was improved when the dough was fermented *L. amylovorus* (Axel et al., 2015), as was also the case when *L. plantarum-based* sourdough was used to produce bread from buckwheat, quinoa, oats, sorghum and teff (Wolter et al., 2014). The results of this study corresponded with the results of (Sanz-Penella et al., 2012) in terms of porosity.

### 3.9. Organoleptic analysis

According to the final scores given to the various breads, there was a significant treatment effect, with the sourdough-based breads ranking above both CO and CA bread. The highest scores were associated with doughs subjected to treatments 8C, 12C and 15C. Each of texture softness, chewiness and taste was significantly affected by starter culture type, and all declined during storage (data not shown). With respect to texture softness, the starter culture type effect was significant in both fresh and stored loaves, with the sourdough-based breads performing better than either the CO or the CA bread. The best treatments with respect to this trait were 7C, 8C, and 9C. With respect to both chewiness and taste, there was a significant effect of starter culture type for stored, but not for fresh loaves. At day zero, treatments 12C and 15C showed the highest score of chewiness, but treatment 16C (CA bread) had the lowest score of chewiness. After four days of storage, doughs subjected to treatments 2C, 5C, 7C, 8C, and 9C scored most highly for chewiness, while CA bread scored poorly. In terms of taste, the CA bread performed worst for both fresh and stored loaves. For the fresh loaves, there was no significant difference between the sourdough-based breads and CO bread, but after four days of storage, breads produced from doughs subjected to treatments 1C, 2C, 3C, 7C, 8C, and 13C scored most favorably. A comparison between the treatments in terms of taste and the amount of diacetyl present suggested that the higher the diacetyl content, the better the taste score (data not shown).

The inclusion of sourdough has a noticeable impact on flavor, in particular, that based on *L. plantarum* and *L. brevis* (Katina et al., 2006). A similar outcome was recorded here for the maize breads. More flavor volatile compounds are formed during the lactic acid fermentation of wholemeal wheat flour-based sourdough than that of white wheat flour-based sourdough (Czerny and Schieberle, 2002). The higher proteolytic activity characteristic of wholemeal flour (Loponen et al., 2004) promotes the accumulation of amino acids such as leucine and proline, which are the flavor precursors generated by standard yeast fermentation and by the Maillard reaction during baking. The generation of some desirable flavor characteristics (overall taste intensity, roasting, and aftertaste) is accompanied by the less desirable flavors such as pungency and staleness, induced by the formation of acetic acid. Acidification is also known to be key for the induction of proteolysis and the enhancement of roasted flavors during dough fermentation (Thiele et al., 2002). Edema (2011), in an evaluation of the effect of *L. plantarum, L. brevis* and *Leuconostoc mesenteroides* on the sensory characteristics of maize-based bread, found that loaves made using a sourdough seeded with all three starter cultures proved to be the most acceptable in terms of taste, texture and overall acceptance. Some sourdough breads were not superior, in terms of their sensory quality, to CO bread and some were even classed as inferior. According to (Crowley et al., 2002), the phenomenon of shrinkage which occurs in sourdough bread results in an increased firmness. During storage, staling occurred gradually and thus breads prepared from all treatments received lower scores. Sourdough breads can develop a pickled taste due to the production of organic acids by the LAB. Note that there was no unanimity as to which of the treatments produced the best tasting bread, reflecting subjective differences between the panel members. (Meignen et al., 2001) have documented that fermentation based on a combination of starter cultures produces a higher number of aromatic compounds than does a single starter culture. In particular, fermentation with *L. brevis* was effective with respect to aromatic compounds, but a greater volume of acetic acid and other aromatic compounds was formed by the combined starter cultures.

### 3.10. Shelf life

In sourdough-based breads, the appearance of mold was delayed compared to CO breads. The high moisture contents of these breads were responsible for their relatively short shelf life. The treatments producing the most mold-resistant breads were 1C, 2C, 5C, 8C, 9C, 10C, 11C, 13C, 14C, and 15C.

## Conclusion

In conclusion, LAB with high EPS production, proteolytic activity, and acidification properties can be considered great for sourdough fermentation. Adding sourdough fermented with LAB reduced the hardness of the resulting bread, its elasticity and the extent of baking loss. The optimal treatments were to use sourdough seeded with *L. brevis* (treatment 2C), with *L. plantarum* (treatment 4C), with *L. brevis* + *L. paralimentarius* (treatment 8C) or with *L. sanfranciscensis* + *L. brevis* + *L. paralimentarius* (treatment 11C). The aforementioned strains, as suitable functional starter cultures for sourdough, could be used as starter cultures in gluten-free sourdough-based breads.

## Acknowledgments

The financial support of the Agricultural Biotechnology Research Institute of Iran (ABRII) [Grant number 3-05-0551-88020] is gratefully acknowledged. The authors declare no conflict of interests and no ethical issues were promulgated.

